# Chromosomal instability accelerates the evolution of resistance to anti-cancer therapies

**DOI:** 10.1101/2020.09.25.314229

**Authors:** Devon A. Lukow, Erin L. Sausville, Pavit Suri, Narendra Kumar Chunduri, Justin Leu, Jude Kendall, Zihua Wang, Zuzana Storchova, Jason M. Sheltzer

## Abstract

Aneuploidy is a ubiquitous feature of human tumors, but the acquisition of aneuploidy is typically detrimental to cellular fitness. To investigate how aneuploidy could contribute to tumor growth, we triggered periods of chromosomal instability (CIN) in human cells and then exposed them to a variety of different culture environments. While chromosomal instability was universally detrimental under normal growth conditions, we discovered that transient CIN reproducibly accelerated the ability of cells to adapt and thrive in the presence of anti-cancer therapeutic agents. Single-cell sequencing revealed that these drug-resistant populations recurrently developed specific whole-chromosome gains and losses. We independently derived one aneuploidy that was frequently recovered in cells exposed to paclitaxel, and we found that this chromosome loss event was sufficient to decrease paclitaxel sensitivity. Finally, we demonstrated that intrinsic levels of CIN correlate with poor responses to a variety of systemic therapies in a collection of patient-derived xenografts. In total, our results show that while chromosomal instability generally antagonizes cell fitness, it also provides phenotypic plasticity to cancer cells that can allow them to adapt to diverse stressful environments. Moreover, our findings suggest that aneuploidy may function as an under-explored cause of therapy failure in human tumors.

## Introduction

One of the most common features of cancer cells is an aneuploid karyotype: approximately 90% of solid tumors and over 50% of hematopoietic cancers display some degree of aneuploidy^1^. Aneuploidy can result from chromosome mis-segregation events during mitosis where one or more chromosomes are not properly segregated to the resulting daughter cells^2^. These events are usually rare in normal cells but occur more frequently during tumor development^3,4^. The increased rate of mis-segregation events commonly found in cancer is called chromosomal instability, or CIN^3,4^.

Despite the ubiquity of chromosomal alterations in human tumors, the induction of aneuploidy generally diminishes a cell’s proliferative capacity^5–7^. By altering the dosage of hundreds or thousands of genes simultaneously, aneuploidy increases a cell’s metabolic requirements and causes widespread problems in protein production, folding, and turnover^8–10^. Additionally, aneuploidy triggers a transcriptional program involving the upregulation of stress response genes and the down regulation of cell cycle genes, often leading to cell cycle arrest and senescence^11–13^. Nonetheless, in human tumors, high levels of aneuploidy are commonly associated with poor patient outcomes, suggesting that aneuploidy could contribute to certain aspects of tumor progression^14–17^.

Recent improvements in cancer therapies have significantly increased the survival and quality-of-life of patients with a diverse range of malignancies^18,19^. However, prolonged patient survival is frequently thwarted by the development of genetic alterations that cause drug resistance, particularly in response to targeted therapies^20^. The standard model of how resistance arises is through the acquisition of mutations that either restore function to the therapeutic target in the presence of the drug or bypass the drug’s effects. For example, in BRAF-driven melanomas, BRAF inhibition commonly results in a therapeutic response, followed by the inevitable outgrowth of tumors harboring secondary mutations in BRAF, NRAS, or MEK1 that allow continued proliferation in the presence of the drug^21^. However, many causes of drug resistance remain unknown: in a recent study of melanoma patients who progressed on BRAF inhibitor treatment, 42% of patients lacked any genetic alteration known to cause BRAFi resistance^22^. Alternate routes to drug resistance that have been identified include cell-state transitions, epigenetic alterations, and the over-expression of drug-efflux pumps, but the relative importance of these pathways remains unknown^23^. In general, our understanding of how cancers develop drug resistance remains incomplete, frustrating our ability to design strategies for long-lasting cancer control.

Studies in yeast have previously linked the acquisition of aneuploidy with decreased sensitivity to a variety of anti-fungal agents^24–26^. For instance, in the pathogenic yeast *Candida albicans*, amplification of a single chromosome causes resistance to the drug fluconazole, and this resistance can be phenocopied by the over-expression of individual genes encoded on this chromosome^27^. We hypothesized that aneuploidy could similarly represent an uncharted cause of drug resistance in human tumors, potentially contributing to the observed association between aneuploidy and poor patient outcomes. To investigate this possibility, we assessed the evolution of drug resistance in cancer cells that had been exposed to periods of chromosomal instability.

### Inhibition of Mps1 accelerates the acquisition of resistance to a BRAF inhibitor

To determine if aneuploidy can drive therapeutic resistance, we sought to transiently induce CIN in cancer cell lines. To achieve this, we treated cells with a small-molecule inhibitor of the spindle-checkpoint kinase Mps1, AZ3146. Treatment with 2µM AZ3146 caused a robust increase in the frequency of mitotic errors in cancer cell lines (Figure S1)^10^. We have previously shown that mutations in the Mps1 kinase domain block the increase in errors caused by AZ3146, verifying that this represents an on-target effect of Mps1 inhibition^17^. We next devised a competition-based strategy to assess if CIN is capable of driving the acquisition of resistance to an anti-cancer therapy (Figure 1A). A BRAF-mutant human melanoma cell line, A375, was transduced to express either of two fluorescent proteins, creating a pair of otherwise-identical cell populations. These fluorescently-labelled populations were then treated for 24-hr with the Mps1 inhibitor (Mps1i) AZ3146 to induce a period of CIN. Next, the drug was washed out, and Mps1i-treated cell populations were co-cultured with an equal number of non-Mps1i-treated cells that had been labeled with a different fluorophore. These cell populations were then passaged for ∼30 days in a variety of growth conditions. At each passage, the relative abundance of each cell population in the coculture was determined by flow cytometry, thus providing a temporal readout of the relative fitness of each population.

**Figure 1.**
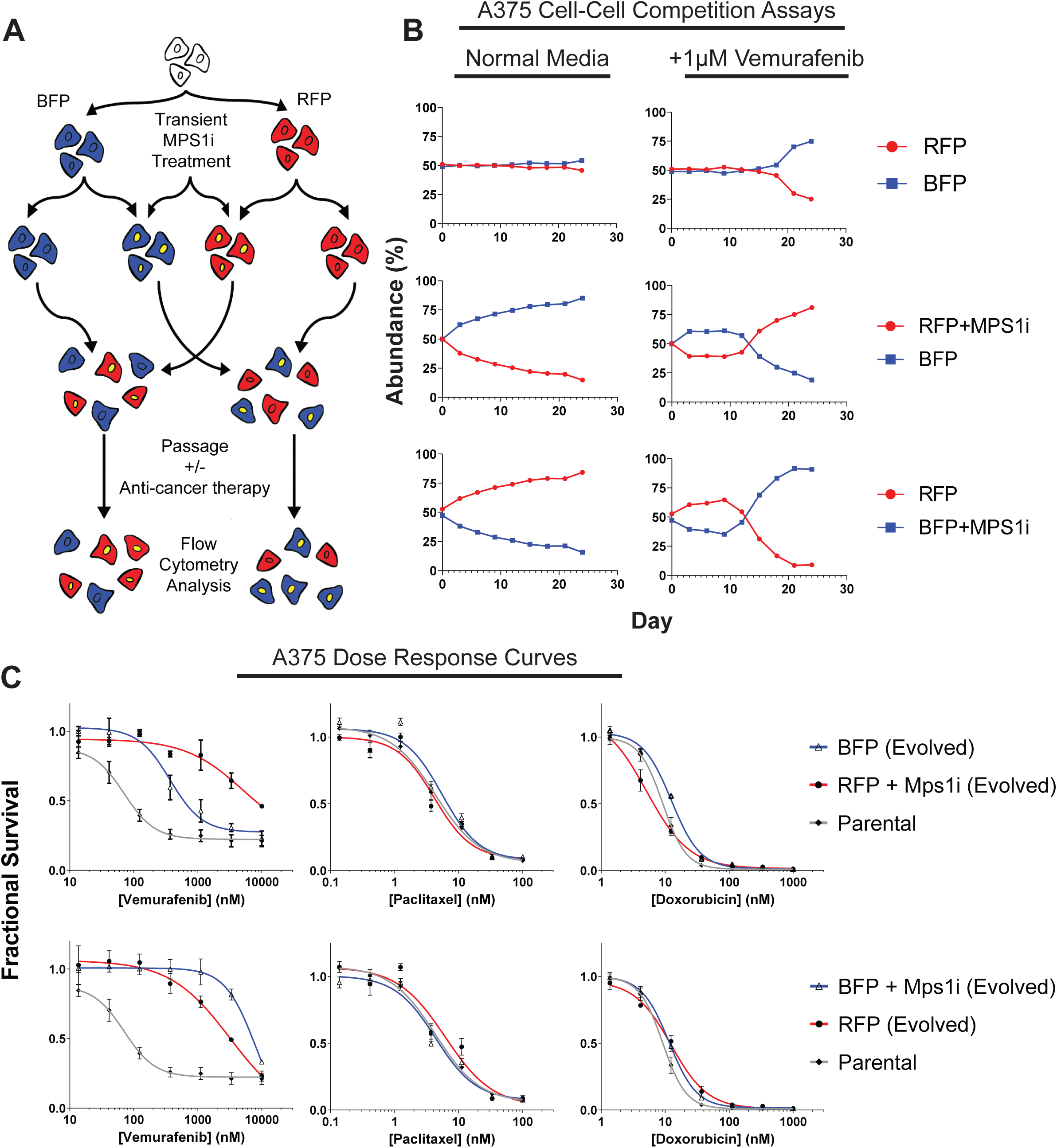
Transient inhibition of Mps1 accelerates the acquisition of resistance to a BRAF inhibitor. (A) Schematic outline of the competition experiment. (B) Relative abundance of each A375 cell population under the indicated growth conditions, as determined by flow cytometry. (C) Dose-response curves of the parental A375 cell line or A375 cells purified by flow cytometry from day 24 of the competition experiments in (B), exposed to the indicated drug.

To assess the effects of CIN on cell fitness, we first conducted competition experiments on melanoma cells grown in standard (drug-free) culture conditions. When BFP+ and RFP+ A375 cells were mixed together at a 50:50 ratio, each population remained at a ∼50% abundance over 24 days in culture, demonstrating that the fluorophores did not significantly affect cell fitness (Figure 1B). However, when either the BFP+ or RFP+ cell populations were pretreated with AZ3146 prior to the competition, the Mps1i-treated population showed a notable decrease in abundance compared to their untreated competitors. For instance, when A375 RFP+ cells were pretreated with AZ3146, they decreased from 50% to 15% abundance over 24 days in culture. These results suggest that periods of chromosomal instability are generally detrimental to cancer cell fitness under standard culture conditions.

We next investigated the effects of CIN on cells exposed to vemurafenib, a BRAF-inhibitor that blocks the growth of BRAF-driven cancer cells like A375. Competitions conducted in media containing 1μM vemurafenib initially resembled the competitions conducted under drug-free conditions, as both the RFP+ and BFP+ populations that were pretreated with AZ3146 decreased in abundance. However, by day 12, both Mps1i-treated populations began to rise in abundance, eventually over-taking the untreated populations and dominating the competitions (Figure 1B). For instance, Mps1-treated RFP+ cells initially decreased from 50% abundance at the start of the competition to 39% abundance at day 9, then increased to 81% abundance by day 24. In the absence of Mps1i-pretreatment, both populations remained at a ∼50:50 frequency for 18 days, and then a slight increase in BFP+ cell fitness was detected (discussed in greater detail below). In total, this experiment suggests that a transient period of CIN can accelerate the ability of melanoma cells to adapt to growth in a BRAF inhibitor.

To further characterize the populations that arose from these competition experiments, BFP+ and RFP+ cells with or without Mps1i-pretreatment were re-isolated by fluorescence activated cell sorting (FACS) and tested in 7-point drug sensitivity assays. All four cell populations that had been passaged in vemurafenib showed a significant decrease in vemurafenib sensitivity compared to their parental line (Figure 1C). However, consistent with the results of the competition assays, cells from the Mps1i-treated populations were more resistant to vemurafenib compared to the populations that were not exposed to Mps1i. These cells were not cross-resistant to two unrelated drugs, paclitaxel and doxorubicin (Figure 1C). This suggests that exposure to an Mps1 inhibitor did not cause a multi-drug resistance phenotype, and that the underlying mechanism(s) of resistance was likely specific to the conditions in which these cells were evolved.

In order to test whether the fitness benefits conferred by exposure to AZ3146 were reproducible, we repeated the above competition experiment in four independent replicates (Figure S2A-D). While the relative abundance of each population varied across replicates, the overall dynamics of the experiment were maintained, and we found that Mps1i-treatment consistently accelerated the acquisition of resistance to BRAF-inhibition (Figure S2A-D). In 9 out of 10 competitions that we conducted, the Mps1i-treated cells eventually dominated the vemurafenib culture, while these same cells never outgrew their competitors in drug-free media. In the absence of Mps1i-treatment, each population was typically maintained at a near-50:50 ratio for several passages, and then one population would eventually increase in abundance. This may reflect selection for underlying genetic or epigenetic alterations that can arise in a heterogeneous cancer cell line and are capable of contributing to drug resistance. However, in contrast to the Mps1i-treated populations, this evolution appears to have occurred randomly, as both RFP+ and BFP+ populations were observed to dominate in independent replicates (e.g., compare Fig. S2C and S2D). Overall, our results demonstrate that a transient period of CIN reproducibly accelerates the acquisition of resistance to vemurafenib, even in heterogeneous populations that harbor other alterations capable of influencing drug resistance.

### Mps1 inhibition can accelerate therapeutic resistance in multiple contexts

We next sought to investigate if transient CIN is capable of accelerating the evolution of resistance to other drugs and in other genetic backgrounds. To test this, we first conducted competition experiments as described in Figure 1A in A375 cells exposed to the microtubule poison paclitaxel. Consistent with our previous results, we found that Mps1i treatment before paclitaxel exposure resulted in a transient loss in cell fitness, followed by the outgrowth of the Mps1i-treated population (Figure 2A). Similar results were obtained in a set of replicate paclitaxel competitions (Figure S3A).

**Figure 2.**
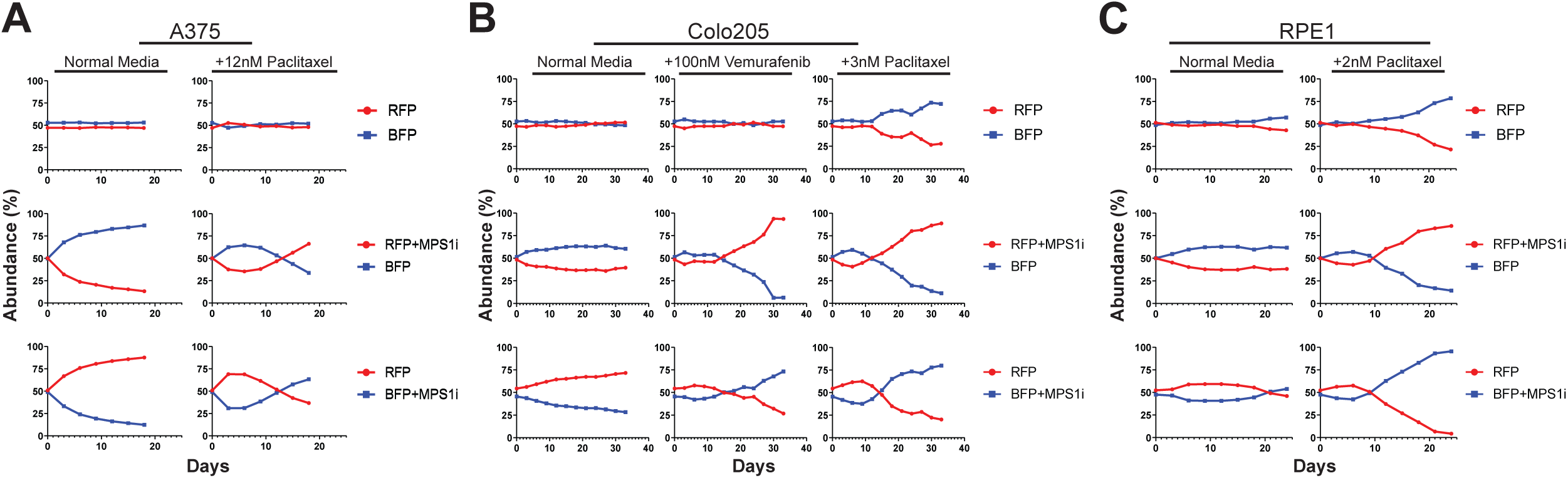
Transient inhibition of Mps1 accelerates the acquisition of drug resistance in multiple genetic backgrounds. (A) Relative abundance of A375 cell populations in normal media or in the presence of paclitaxel as determined by flow cytometry. (B) Relative abundance of Colo205 cell populations in normal media, in the presence of vemurafenib, or in the presence of paclitaxel, as determined by flow cytometry. (C) Relative abundance of RPE1 cell populations in normal media or in the presence of paclitaxel as determined by flow cytometry.

Next, we investigated whether this phenomenon was exclusive to A375 cells. To test this, we performed cellular competitions in a BRAF-mutant colorectal cancer cell line, Colo205. Under drug-free growth conditions, Mps1i-treated cell populations decreased in abundance from ∼50% to ∼30% when competed for 30 days against untreated cells (Figure 2B). However, when competitions were performed in either vemurafenib or paclitaxel, we observed that the Mps1i-treated cell populations reproducibly dominated each culture. For instance, when Colo205 cells were competed in 100nM vemurafenib, Mps1i-treated populations initially dropped to ∼40% abundance, then increased in abundance from day 15 to day 33, at which point the Mps1i-treated populations made up ∼85% of their cultures. Similar results were obtained in a set of replicate competitions in vemurafenib, though the magnitude of the benefit conferred by Mps1i treatment varied between replicates, underscoring the stochastic nature of these evolutionary adaptations (Figure S3B).

To explore whether the benefits conferred by transient CIN were restricted to malignant cells, we next examined the consequences of Mps1 exposure in immortalized retinal pigment epithelial (RPE1) cells. As we observed in A375 and Colo205 cancer cells, Mps1i pretreatment strikingly accelerated the growth of RPE1 cells cultured in paclitaxel in multiple independent competitions (Figure 2C and Figure S3C). Interestingly, the benefits of transient Mps1i treatment were not universal: exposure to AZ3146 did not increase the fitness of EGFR-driven PC9 lung cells cultured in the EGFR inhibitor gefitinib or of BRAF-driven SK-MEL-5 cells cultured in vemurafenib (Figure S4; discussed in greater detail below). In total, these results indicate that exposure to an Mps1 inhibitor can accelerate the evolution of resistance to multiple targeted and cytotoxic agents in a variety of genetic backgrounds.

### Cells that acquire drug resistance after Mps1i treatment display specific recurrent aneuploidies

To explain the dynamics that we observed in our cellular competition experiments, we hypothesized that treatment with an Mps1 inhibitor creates an initial population of cells with diverse karyotypes. As most of these aneuploidies are detrimental to cell fitness^28,29^, the Mps1i-treated populations are initially outcompeted by the untreated populations. However, certain rare aneuploidies could promote resistance to specific drugs, as has previously been observed in yeast cells treated with antimycotics^24,26,30^. Over time, growth in the presence of anti-cancer drugs could select for cells harboring aneuploidies that are advantageous under those specific conditions, revealing a beneficial role for CIN in the evolution of drug resistance (Figure 3A).

**Figure 3.**
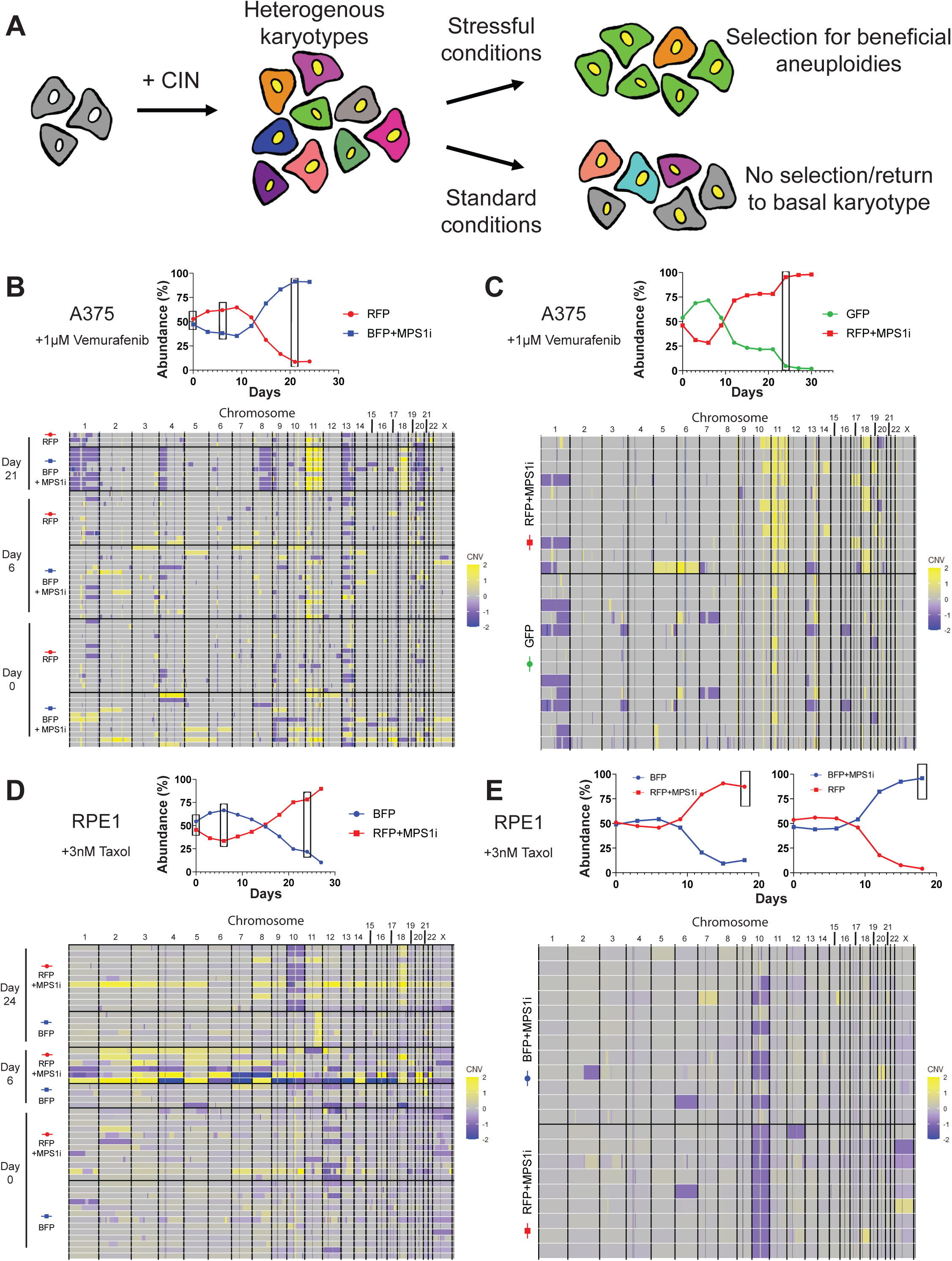
Cells that develop drug resistance following Mps1i exposure display recurrent aneuploidies. (A) Diagram modeling selection for beneficial aneuploidies under stressful conditions. (B) (Top) Plot displaying the competition of A375 cells in vemurafenib from which cells were isolated for single cell sequencing. Boxes indicate time points when cells were isolated. (Bottom) Heatmap displaying the karyotypic alterations of single cells isolated from various time points of the above competition. (C) (Top) Plot displaying the competition of A375 cells in vemurafenib from which cells were isolated for single cell sequencing. Box indicates time point when cells were isolated. (Bottom) Heatmap displaying the karyotypic alterations of single cells isolated from the above competition. (D) (Top) Plot displaying the competition of RPE1 cells in paclitaxel from which cells were isolated for single cell sequencing. Boxes indicate time points when cells were isolated. (Bottom) Heatmap displaying the karyotypic alterations of single cells isolated from various time points of the above competition. (E) (Top) Plot displaying the competition of RPE1 cells in paclitaxel from which cells were isolated for single cell sequencing. Box indicates time point when cells were isolated. (Bottom) Heatmap displaying the karyotypic alterations of single cells isolated from the above competition. N.B. – Competition plots shown in (B), (C), and (E) were previously shown in Figures 1B, S2A, and S3C, respectively.

To investigate this hypothesis, we performed single-cell genomic sequencing on cells from different competition experiments. First, we isolated single A375 cells from either Mps1i-treated (BFP-expressing) or untreated (RFP-expressing) cells competed in vemurafenib for 0, 6, or 21 days. We performed read-depth analysis to identify copy number changes in each cell relative to the untreated parental cell population. As expected, the Mps1i pulse caused a significant increase in karyotype heterogeneity that was apparent at day 0 and day 6 relative to the untreated cells (Figure 3B). Interestingly, at day 21, when the Mps1i-treated population had overtaken the untreated population, the heterogeneity among Mps1i-treated cells had declined, and a nearly-clonal aneuploid karyotype had emerged. These new aneuploidies included a gain of chromosome 11 (detected in 7/8 cells), a gain of chromosome 18 (detected in 7/8 cells), a loss of 40 Mb from chromosome 4p (detected in 8/8 cells), and several other alterations. These karyotypic changes were either not detected or were recovered at a lower frequency in the untreated RFP-expressing population. If these aneuploidies conferred a fitness advantage on A375 cells grown in vemurafenib, then we would anticipate that they would independently recur in separate competitions. To test this, we performed single-cell genomic sequencing from cells collected at day 27 in a distinct A375+vemurafenib competition. In this assay, we again recovered amplifications of chromosome 11 (detected in 9/11 cells) and chromosome 18 (detected in 5/11 cells), but the loss of chromosome 4p was not observed (Figure 3C). We conclude that certain aneuploidies are strongly associated with vemurafenib resistance in A375 cells. Other aneuploidies that we observed could conceivably represent “passenger” alterations, particularly if they first arose in cells that also harbored gains of chromosome 11 and/or 18.

We next sought to discover whether the evolutionary pressure conferred by anti-cancer agents selects for clonal aneuploidies in other cell lines. To address this question, we performed single-cell sequencing on three independent competitions of Colo205 cells cultured in vemurafenib. We observed that Mps1i-treated Colo205 cells nearly always acquired gains of chromosome 7 when grown in vemurafenib, as this aneuploidy was observed in 24/38, 27/32, and 29/32 cells in the three independent competitions (Figure S5A-C). In contrast, amplifications of chromosome 11 and 18 were rarely recovered in these experiments. This suggests that the same anti-cancer drug can select for different aneuploidies in different genetic backgrounds, and no universal “vemurafenib-resistance” karyotype exists.

Finally, we analyzed karyotypes from Mps1i-treated and untreated RPE1 cells competed in paclitaxel. We isolated single cells from either Mps1i-treated (RFP-expressing) and untreated (BFP-expressing) cells competed in 3nM paclitaxel for 0, 6, and 24 days. At day 0 and 6, an increase in karyotype heterogeneity is apparent in the Mps1i-treated cells compared to the untreated counterparts (Figure 3D). By day 24, when the Mps1i-treated population dominated the competition, their karyotypic heterogeneity had declined and a near clonal karyotype emerged. The dominant Mps1i-treated cells displayed loss of chromosome 10 (detected in 8/11 cells) and gain of chromosome 18 (detected in 8/11 cells). Single-cell analysis from an independent experiment again revealed chromosome 10 loss (detected in 9/9 RFP+Mps1i cells and 5/12 BFP+Mps1i cells), suggesting that this karyotypic alteration drove resistance to paclitaxel (Figure 3D). In total, we conclude that transient chromosomal instability can generate populations of karyotypically-heterogeneous cells, and certain aneuploidies reproducibly arise to near-clonal levels when cells are competed under the selective pressure of anti-cancer drugs (Table 1).

**Table 1.** Summary of Single-Cell Karyotypic Analysis of Mps1i-Treated Cell Lines.

| Cell Line | Drug | Mps1i-related Resistance | Recurrent Aneuploidies<sup>a</sup> | Sporadic Aneuploidies<sup>b</sup> |
| --- | --- | --- | --- | --- |
| A375 | Vemurafenib | Yes | Gain: 11, 18 | Loss: 1, 4p, 8q, 13, 20 |
| RPE1 | Paclitaxel | Yes | Loss: 10 | Gain: 18 |
| Colo205 | Vemurafenib | Yes | Gain: 7 | Gain: 8, 18<br>Loss: 1q, 13 |
| PC9 | Gefitinib | No | Whole-genome duplication | N/A |

<sup>a</sup> Aneuploidies were considered recurrent if they were present in at least 50% of Mps1i-treated cells in two or more independent replicates
<sup>b</sup> Aneuploidies were considered sporadic if they were present in at least 50% of Mps1i-treated cells in only one replicate

In order to better understand why Mps1i-treatment failed to promote the acquisition of drug resistance in certain contexts, such as PC9 cells competed in gefitinib, we performed single-cell sequencing on cells isolated from this experiment (Figure S6). Interestingly, we observed that the Mps1i-treated PC9 cells from two independent competitions often displayed whole-genome duplications, while such events were rare in other cell types. This observation is consistent with the results of our live-cell imaging analysis, which revealed that AZ3146 induced mitotic slippage at a much higher rate in PC9 cells compared to A375 cells (39% vs. 17%; Figure S1B). Thus, AZ3146 exposure may cause tetraploidization events through mitotic slippage in PC9 cells, and this type of alteration may be less likely to directly drive drug resistance. Additionally, in the PC9 cells that did not exhibit evidence of whole-genome duplications, we did not observe enrichment of any individual aneuploidy. Thus, it is possible that the level of single-chromosome missegregation events generated by this treatment did not produce enough karyotypic diversity to confer drug resistance, or that other, aneuploidy-independent resistance mechanisms are more common in this genetic context.

### A recurrent aneuploidy observed in cellular competition experiments is sufficient to confer resistance to paclitaxel

We hypothesized that many of our single-cell sequencing experiments recovered the same aneuploidies in independent competitions because these aneuploidies decreased a cell’s sensitivity to the anti-cancer agent that was being applied. Alternately, it is possible that these aneuploidies are “passenger” events, and their prevalence during cellular competitions occurs by chance. To differentiate between these two possibilities, we sought to discover whether an aneuploidy that we detected in our single-cell sequencing experiments could in fact confer drug resistance.

To accomplish this, we isolated and karyotyped clones derived from single RPE1 cells. We discovered that untreated RPE1 clones occasionally displayed whole-chromosome aneuploidies, but monosomies were generally unstable and conferred a severe fitness disadvantage. However, deleting the tumor suppressor p53 from RPE1 cells allowed us to recover certain monosomies and conduct short-term experiments on them (Chunduri et al., manuscript in revision). We successfully isolated one RPE1 p53-/- clone that had spontaneously lost a copy of chromosome 10, which was the most common aneuploidy that we observed in RPE1 cells cultured in paclitaxel (Figure 4A-B). (Note that the parental RPE1 cell line is disomic for chromosome 10p and trisomic for chromosome 10q, and thus these clones are monosomic for 10p and disomic for 10q). As a control, we also isolated an RPE1 clone that had lost a copy of chromosome 13, which we never observed in our paclitaxel-resistance experiments. We then competed these chromosome-loss clones against a parental population of RPE1 p53-/- cells. We found that losing chromosome 13 had minimal effect on cellular fitness in either normal media or in media that contained paclitaxel (Figure 4C). Losing chromosome 10 reduced the fitness of RPE1 cells in normal media, as this clone decreased to ∼12% abundance over 11 days in culture. Remarkably, this same aneuploidy increased in abundance from 46% to 89% when competed in media containing paclitaxel (Figure 4C). These results demonstrate that an aneuploidy that compromises proliferation under normal growth conditions can enhance cell fitness in the presence of a chemotherapy agent. Moreover, these findings indicate that the aneuploidies recovered in our cellular competition assays are in fact capable of directly causing drug resistance.

**Figure 4.**
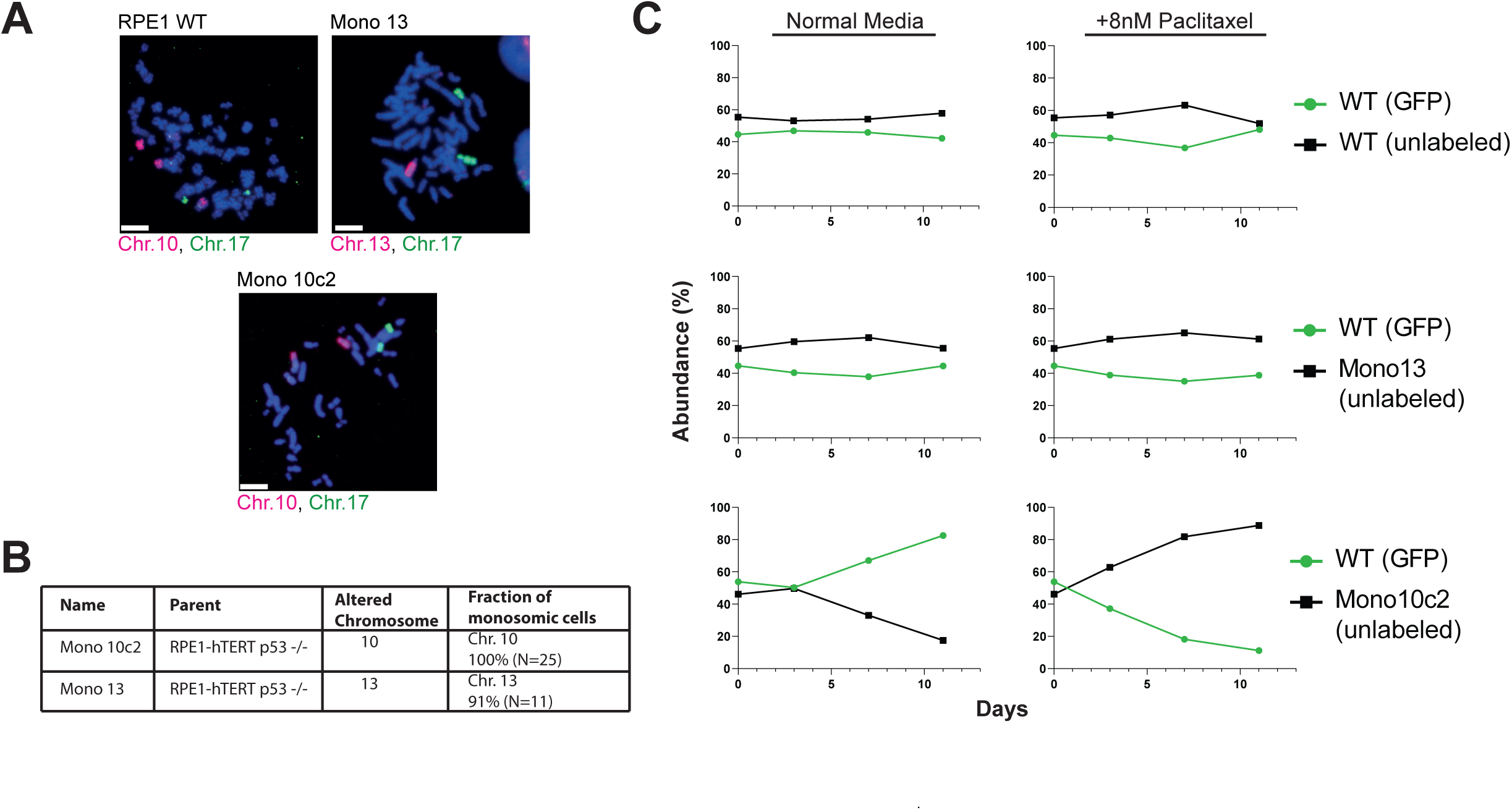
A recurrent aneuploidy recovered in cellular competition experiments is sufficient to confer resistance to paclitaxel. (A) Micrographs of chromosome paints for chromosomes 10, 13, and 17 in RPE1 clones to confirm monosomies. (B) Summary of the chromosome painting experiments. (C) Relative abundance of GFP-expressing RPE1 cells competed against unlabeled monosomies under normal growth conditions or in the presence of paclitaxel.

### High levels of chromosomal instability correlate with treatment resistance in patient-derived xenografts

Thus far, we have reported that CIN caused by an Mps1 inhibitor accelerates the evolution of resistance to anti-cancer therapies. We next sought to discover whether endogenous levels of CIN, in the absence of a CIN-inducing agent, could similarly interfere with treatment efficacy. To investigate this question, we analyzed a large dataset of patient-derived xenografts (PDXs) that had been profiled over multiple passages in mice^31^. To determine the level of endogenous CIN in each PDX, we calculated the average number of arm-length copy-number changes that occurred each time a PDX was passaged. We defined PDXs that exhibited fewer than one arm-length change per passage as “low CIN” cancers, while PDXs that exhibited four or more arm-length changes per passage were defined as “high CIN” cancers.

Mice bearing these PDXs were treated with a variety of anti-cancer agents and subsequent changes in tumor volume were determined. RECIST criteria was used to identify tumors that exhibited a therapeutic response, stable disease, or progressive disease, during systemic treatment with each of several different drugs or drug combinations^31^. We discovered that, across all cancer types and therapies, 11% of low CIN PDXs exhibited a partial or complete therapeutic response to drug treatment, compared to only 3% of high CIN cancers (Figure 5A; P < .002, Fisher’s exact test). Likewise, 69% of high CIN cancers exhibited progressive disease upon treatment, compared to 58% of low CIN cancers (Figure 5B; P < .03, Fisher’s exact test). Similar overall trends were observed in individual cancer types. For instance, 10% of low CIN breast cancer PDXs exhibited a therapeutic response to the given treatments, while 0% of high CIN breast cancers responded (Figure 5A; P < .007. Fisher’s exact test).

**Figure 5.**
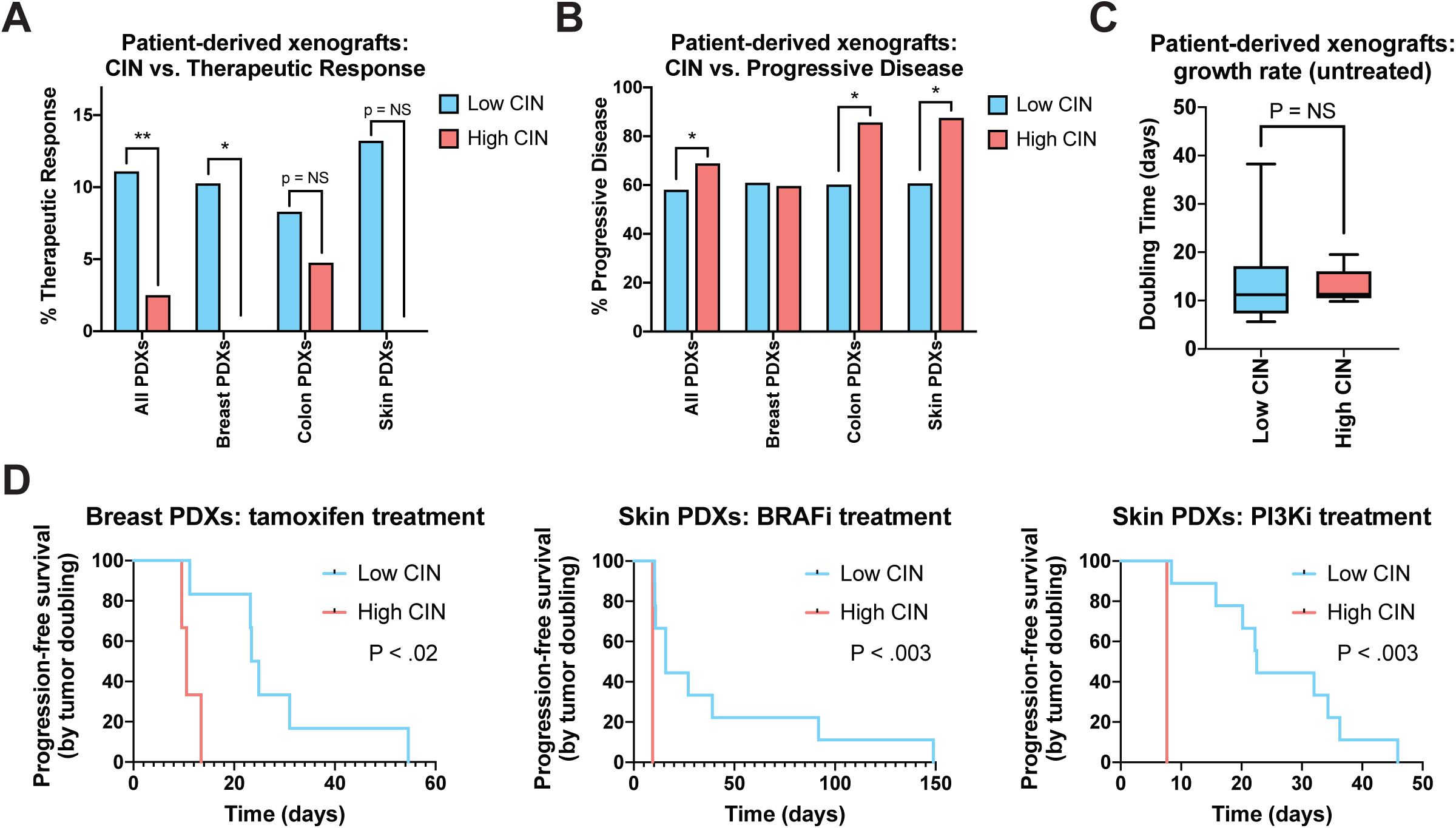
High levels of endogenous CIN are associated with poor drug responses in a set of patient-derived xenografts. (A) The percent of PDXs that displayed a partial or complete therapeutic response to systemic therapy, sorted according to their degree of CIN. (B) The percent of PDXs that displayed progressive disease in response to systemic therapy, sorted according to their degree of CIN. (C) Box-plots showing the time required for an untreated PDX to double in volume, sorted according to their degree of CIN. Boxes indicate the 25^th^, 50^th^, and 75^th^ percentiles, while the bars represent the 10^th^ and 90^th^ percentiles. (D) Kaplan-Meier survival analysis of progression-free survival, defined as the time required for a treated PDX to double in volume, for PDXs treated with the indicated therapies.

We considered the possibility that high CIN cancers grew more rapidly than low CIN cancers, thereby explaining this difference in therapeutic response rates. However, in mice that were not treated with any anti-cancer agents, both low CIN and high CIN PDXs exhibited a median tumor-volume doubling time of 11 days, suggesting that they exhibited similar proliferative capacities (Figure 5C). Finally, we sought to determine whether different response patterns could be detected in PDXs treated with specific therapeutic agents. We calculated a surrogate “progression-free survival” period for each PDX, based on the time required for the tumor volume to double during treatment. Consistent with the *in vitro* competition experiments that we conducted, we discovered that high CIN skin cancer PDXs treated with BRAF inhibitors exhibited a significantly shorter progression-free survival time compared to low CIN skin cancer PDXs (Figure 5D, P < .003, log-rank test). Similar results were observed in several other treatment conditions, including breast cancer PDXs treated with tamoxifen and skin cancer PDXs treated with PI3K inhibitors (Figure 5E-F). We conclude that high levels of endogenous CIN are correlated with an increased likelihood of progressive disease and therapeutic resistance in a variety of human cancer types.

## Discussion

Here we demonstrated that artificially induced CIN through inhibition of the SAC kinase Mps1 can accelerate the acquisition of resistance to both targeted and cytotoxic therapies. Single cell karyotypic analysis of the drug-resistant cells across multiple replicates showed consistent but unique karyotypic alterations for each drug-cell line combination. Contemporaneously, Ippolito *et al*. employed a similar experimental approach to study the role of CIN in promoting drug resistance. Using a different inhibitor of Mps1, different cell lines, and different anti-cancer therapies, they also report that CIN is capable of promoting drug resistance. These findings highlight the generalizability of this phenomenon as a novel cause of anti-cancer therapy failures.

While CIN and the aneuploidies that result from CIN cause numerous cellular stresses, chromosomal errors are ubiquitous during tumorigenesis, and the level of aneuploidy tends to be highest in aggressive cancers associated with poor patient outcomes^2,6,7,17,32^. It remains unclear to what extent the aneuploidy that arises during tumorigenesis functions as a driver of cancer progression, and to what extent aneuploidy occurs simply as a byproduct of the loss of checkpoint control that often occurs in advanced malignancies. Previous studies have shown that CIN can accelerate the adaptation to oncogene withdrawal in mouse models of KRAS and BRAF driven cancers, which often displayed recurrent aneuploidies^33,34^. Similarly, our work shows that CIN can drive the evolution of drug-resistant cells with recurrent aneuploidies, and it demonstrates that one such aneuploidy is sufficient to confer drug resistance. We believe that our findings coupled with those of Ippolito *et al*. may represent a novel mechanistic category of drug resistance that could help explain the large fraction of tumors for which no known resistance mechanism can be identified. The model presented in this paper proposes a causative role for CIN and aneuploidy in promoting drug resistance through the generation of karyotypic heterogeneity upon which selection can act. CIN allows cancer cells to sample the fitness landscape by acquiring random aneuploidies, most of which will be deleterious, and a few of which will provide a fitness benefit under selective conditions. Over the course of tumor progression, this selection for beneficial aneuploidies arising from CIN would result in higher degrees of aneuploidy, causing the association between CIN, aneuploidy, and aggressive malignancies.

It has previously been reported that cancer cell lines with CIN show a reduced sensitivity to a broad range of cytotoxic and targeted therapies compared to their euploid counterparts^35^. However, we observed that the immediate consequence of CIN is reduced fitness under both normal and stressful growth conditions. We only detected an increase in the fitness of CIN cells after several passages, and single-cell sequencing revealed that this increase co-occurred along with the rise of certain clonal aneuploidies. This suggests that CIN does not provide “intrinsic” resistance to anti-cancer therapies, and instead endows a genomic plasticity to cancer cells that allows them to acquire beneficial aneuploidies. In support of this model, it has previously been observed that certain karyotypic alterations are associated with improved proliferation in serum-starved or hypoxic conditions, underscoring aneuploidy’s ability to promote growth in stressful environments^36^.

In our experiments, we recovered the same aneuploidies in independent competitions, suggesting that these chromosomal alterations drove the drug-resistance phenotype. Moreover, by reconstituting one aneuploidy in drug-naïve cells, we were able to demonstrate that this karyotypic change was sufficient to decrease paclitaxel sensitivity. Aneuploidy has been proposed to promote tumor development by increasing the dosage of growth promoting oncogenes and decreasing the dosage of growth inhibitory tumor suppressors^37–39^. Similarly, we hypothesize that the altered expression of certain dosage-sensitive genes encoded on the chromosomes we recovered drives selection for these specific aneuploidies. However, the identity of these gene(s) is at present unknown. Chromosome-specific and drug-specific interactions may complement the multi-drug resistance phenotype that has been observed in near-tetraploid cells that have undergone whole-genome duplications^40–42^. However, Mps1i treatment did not accelerate evolution in every context tested. We speculate that different classes of mitotic errors may exhibit different effects on cellular evolvability, and in certain contexts whole-genome duplications, as observed in PC9 lung cancer cells, may promote senescence rather than drug resistance.

Finally, in this work, we applied pharmacological inhibitors of Mps1 in order to generate periods of chromosomal instability. Mps1 inhibitors are also being investigated as potential cancer therapies, and at least five different Mps1-targeting compounds have entered clinical trials^43^. However, our results underscore how CIN can drive genomic plasticity in cancer and promote the development of drug resistance in a variety of contexts. These findings suggest that Mps1 inhibitors should be used cautiously in the clinic, so as not to inadvertently accelerate the evolution of drug resistance in cancer patients.

## Methods

### Cell lines and tissue culture

The identity of each cell line was verified by STR profiling (University of Arizona Genetics Core, Tucson, AZ). A375, RPE1, SK-MEL-5, and SK-MEL-28 cell lines were cultured in DMEM (Gibco) with 10% FBS, 2mM glutamine, and 100 U/mL penicillin and streptomycin. Colo205 and PC9 cell lines were cultured in RPMI-1640 (Gibco) with 10% FBS, 2mM glutamine, and 100 U/mL penicillin and streptomycin. All cell lines were grown in a humidified environment at 37°C and 5% CO_2_ and passaged every 3 days.

### Virus generation and transduction

Retrovirus for constructs expressing EGFP (Addgene #160226), EBFP (Addgene #160227), and mRFP1 (Addgene #160228) were generated using calcium phosphate transfection previously described^44^. Virus was collected 48-hours and 72-hours post transfection and filtered through a 0.45μm syringe. Collected virus was either immediately applied to target cells with 4μg/mL polybrene for transduction or stored at −80°C for later use.

### Live cell imaging

A375 and PC9 cells harboring an GFP-H2B construct (Addgene #26790) were plated on 8-well μ-slides (Ibidi; Cat. No. 80826) and half were treated with 2μM AZ-3146 (Selleck Chemicals; Cat. No. S2731) 3-hours prior to imaging. Live-cell imaging was performed in a humidified environment at 37°C using a spinning-disc confocal microscopy system (UltraVIEW Vox; PerkinElmer) and a charge-coupled device camera (ORCA-R2; Hamamatsu Photonics) fitted to an inverted microscope (DMI6000 B; Leica) equipped with a motorized piezo-electric stage (Applied Scientific Instrumentation). Overnight imaging was performed using a Plan Apochromat 20X 0.7 NA air objective with camera binning set to 2×2. Image acquisition and analysis were performed using Volocity version 6.3 (PerkinElmer). Cells undergoing mitosis were tracked from nuclear envelope breakdown to anaphase exit, during which time errors such as lagging chromosomes, polar chromosomes, multipolar mitoses, anaphase bridges, and mitotic arrest were observed and scored.

### Competition experiments

Cell lines transduced with different fluorophores were cultured in normal growth conditions or pulse-treated with 2μM AZ-3146 for the duration of approximately two cell doubling times (24 hours for A375 and RPE1; 48 hours for Colo205, PC-9, SKMEL-5). Pulse-treated cells were then given a three-day recovery passage in normal growth conditions. Pulse-treated cells and non-treated cells harboring a different fluorophore were then mixed in a 50:50 ratio confirmed by flow cytometry (MACSQuant; Miltenyi Biotec). Gates were established using non-fluorescent parental cell lines as a negative control. Cell mixtures were then cultured in normal growth media or media supplemented with anti-cancer therapeutics. Cell mixtures were passaged every three days and the relative abundance measured by flow cytometry.

### Single-cell sequencing and SMASH sequencing

The single-cell sequencing protocol was adapted from Baslan,*et al*^45^. Cells from competition experiments were single-cell sorted into 96-well PCR plates (ThermoFisher Scientific; Cat. No. AB0731) containing lysis buffer (0.1% SDS, 2% Triton) and incubated at 65°C for 1 hour. Genomic DNA was enzymatically digested with NlaIII (NEB; Cat. No. R0125). Both ends of genomic fragments were tagged using oligonucleotides containing a cell-barcode, a universal primer, and several random nucleotides (varietal tags) through ligation and extension reactions. Cell barcodes allow for multiplexing of the samples and the varietal tags allow for counting of unique initial DNA fragments. After amplification with a universal primer, fragment size selection was performed with Agencourt AMPure XP beads (Beckman Coulter; A63881) to obtain fragments appropriate for sequencing. Barcoded sequencing adaptors were ligated to the ends of the fragments to allow for further multiplexing and to prepare the fragments for next generation sequencing. Next generation sequencing was performed on an Illumina MiSeq. Reads were demultiplexed and used to identify karyotypes as described in ref^46^. Heatmaps of single-cell karyotypes compared to parental basal karyotypes were generated in R.

Basal karyotypes of each cell line were identified through SMASH sequencing (Wang et al., 2016). Cells were trypsinized, resuspended in PBS, and pelleted. Total Genomic DNA was then extracted from the cell pellets via Qiagen QIAamp kit (Cat. No. 51036). The genomic DNA was then fragmented with dsDNA Fragmentase (NEB, Cat. No. M0348L) to a mean size of ∼40bp followed by random ligation to form chimeric fragments approximately 400-700bp in length. Size selection of suitably sized fragments for NGS library preparation was performed with Agencourt AMPure XP beads (Beckman Coulter, Cat. No. A63881). Illumina compatible NEBNext Multiplex Dual Index Primer pairs and Adaptors (New England Biolabs, Cat. No. E6440S) were then ligated to the ends of the size-selected chimeric fragments. These fragments were then sequenced on an Illumina MiSeq. NGS-generated reads were demultiplexed and mapped to generate karyotypes as described in ref^46^.

### Generation of monosomies and chromosome paints

The monosomic cell lines arose spontaneously following depletion of TP53 gene from human retinal pigment epithelium cell line RPE1 immortalized with hTERT overexpression. Briefly, the gRNA against *TP53* was cloned in pX330 vector (Addgene: 42230) according to a modified protocol from Shalem, *et al* 2014 ^47^ and used to transfect RPE WT cells. Single cell derived clones were verified for the loss of p53 expression by sensitivity to Nutlin and immunoblotting for p53 and p21. The copy number status of the single cell derived clones were verified by low-pass whole genome sequencing. All the cell lines were cultured in DMEM + GlutaMAX™-I medium (Gibco) supplemented with 10% FBS and penicillin and streptomycin at 37°C in a humidified 5% CO_2_ incubator. To minimize the occurrence of secondary genomic changes, original stocks were thawed for every experiment and maintained for maximum of 4 to 5 passages.

Additional validation of monosomies was performed with chromosome paints. Cells were treated with 400 ng/mL colchicine for 5-6 h, trypsinized, and pelleted. Cell pellets were resuspended in 75 mM KCl and incubated for 10–15 min at 37°C. Cells were pelleted at 1000 rpm for 10 min and suspended in 3:1 methanol/acetic acid to fix the cells, then washed several times in 3:1 methanol/acetic acid. Fixed cells were dropped on a glass slide and dried at room temperature for 15 min. Each sample was labeled with chromosome FISH probes (Chrombios) specific for a monosomic chromosome and a control chromosome as per manufacturer’s instructions. Briefly, chromosome spreads were incubated with probe mixture (1 μL of each probe, adjusted to 10 μL with HybMix buffer). After denaturation at 72°C for 6 min, slides were kept at 37°C in a humid chamber overnight. Slides were washed for 5 min in 2x saline sodium citrate (SSC) solution and then for 1 min in prewarmed 70°C 0.4X SSC, 0.1% Tween solution, and, finally, in 4x SSC, 0.1% Tween solution for 5 min at room temperature. Then slides were incubated for 30 min at 37°C with 100 μL fluorescein isothiocyanate (FITC) mouse anti-digoxin (Jackson Immuno Research) solution (1:300 in 4X SSC/0.1% Tween) and washed twice in 45°C pre-warmed 4x SSC/0.1% Tween solution for 5–10 min. Finally, cells were stained with DAPI and microscopic analysis was carried out using 3i software and spinning disc confocal microscopy (see below). For each sample, at least 25 metaphases were captured and analyzed (Chunduri et al., manuscript in preparation).

### Analysis of PDX models

Data on patient-derived xenograft drug responses was acquired from Gao et al^31^. The degree of chromosomal instability in each model was determined as described in Ben-David et al^48^. Low CIN PDXs were defined as PDXs that displayed an average of less than one arm-length copy number change per passage. High CIN PDXs were defined as PDXs that displayed an average of greater than four arm-length copy number changes per passage. Therapeutic responses were determined according to RECIST criteria, as described in Gao^31^. Progression-free survival times were defined as the time required for a treated tumor to double in volume.

## Acknowledgments

Research in the Sheltzer Lab is supported by an NIH Early Independence award (1DP5OD021385), NIH grant R01CA237652-01, Department of Defense grant W81XWH-20-1-068, a Damon Runyon-Rachleff Innovation award, an American Cancer Society Research Scholar Grant, and a grant from the New York Community Trust. This work was also supported by grants to M. Wigler from the Simons Foundation, Life Sciences Founders Directed Giving-Research (award numbers 519054) and the Breast Cancer Research Foundation (award number 19-174). We thank Uri Ben-David (Tel Aviv University) for assistance with PDX analysis.

This work was performed with assistance from CSHL Shared Resources, including the CSHL Flow Cytometry Shared Resource, which are supported by the Cancer Center Support Grant 5P30CA045508.

## LEGEND

**Figure S1.**
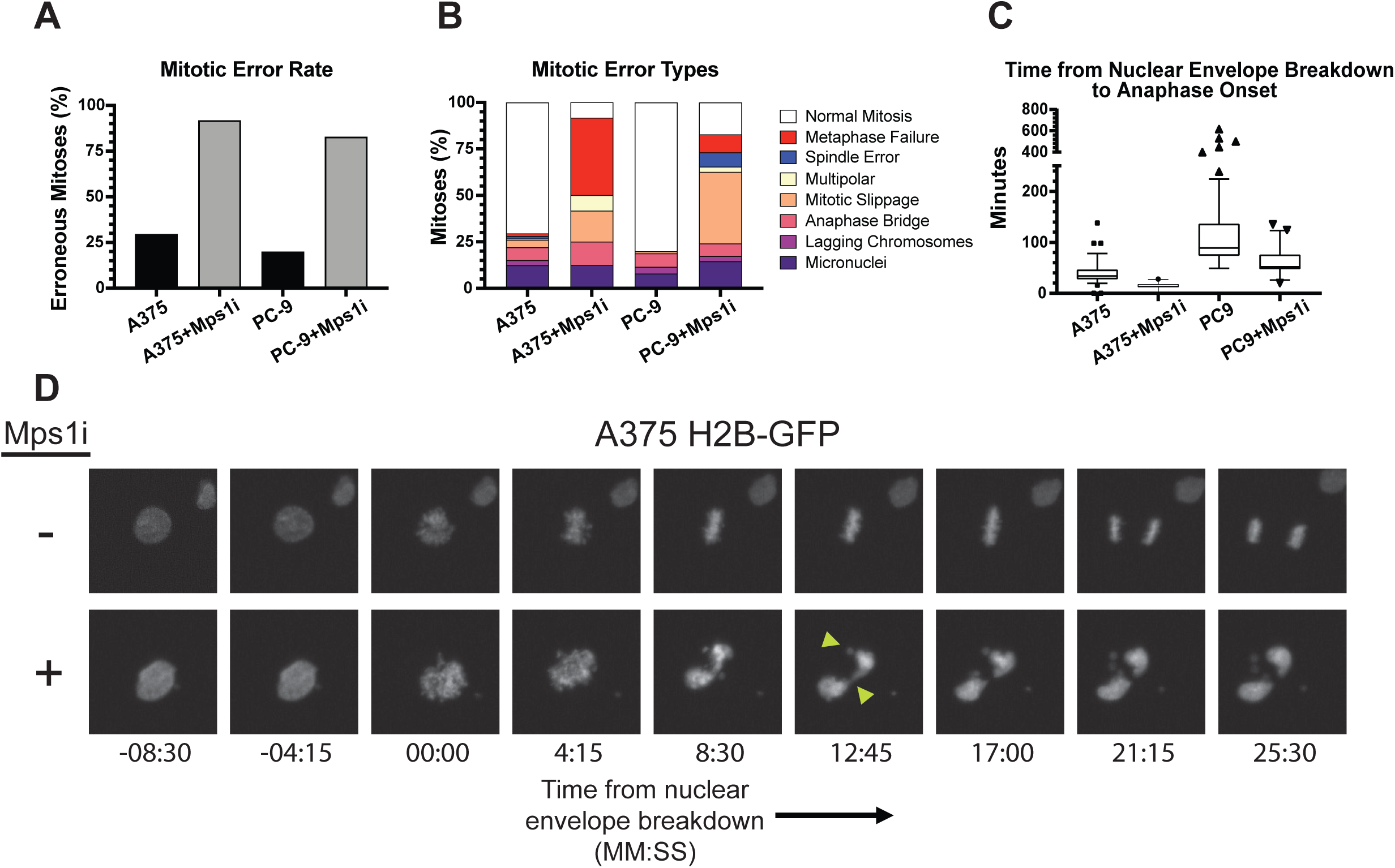
Induction of chromosomal instability by Mps1 inhibitor treatment. Related to Figure 1. (A) Bar graph of the percent of erroneous mitoses in GFP-H2B A375 and PC-9 cells in the presence or absence of the Mps1 inhibitor AZ3146. (B) Bar graph of the percent of mitoses that display various chromosome missegregation events in the presence or absence of AZ3146. (C) Tukey plot of the duration of time cells spent in mitosis in the presence or absence of AZ3146. (D) Image series of representative mitoses from cells in the presence or absence of AZ3146.

**Figure S2.**
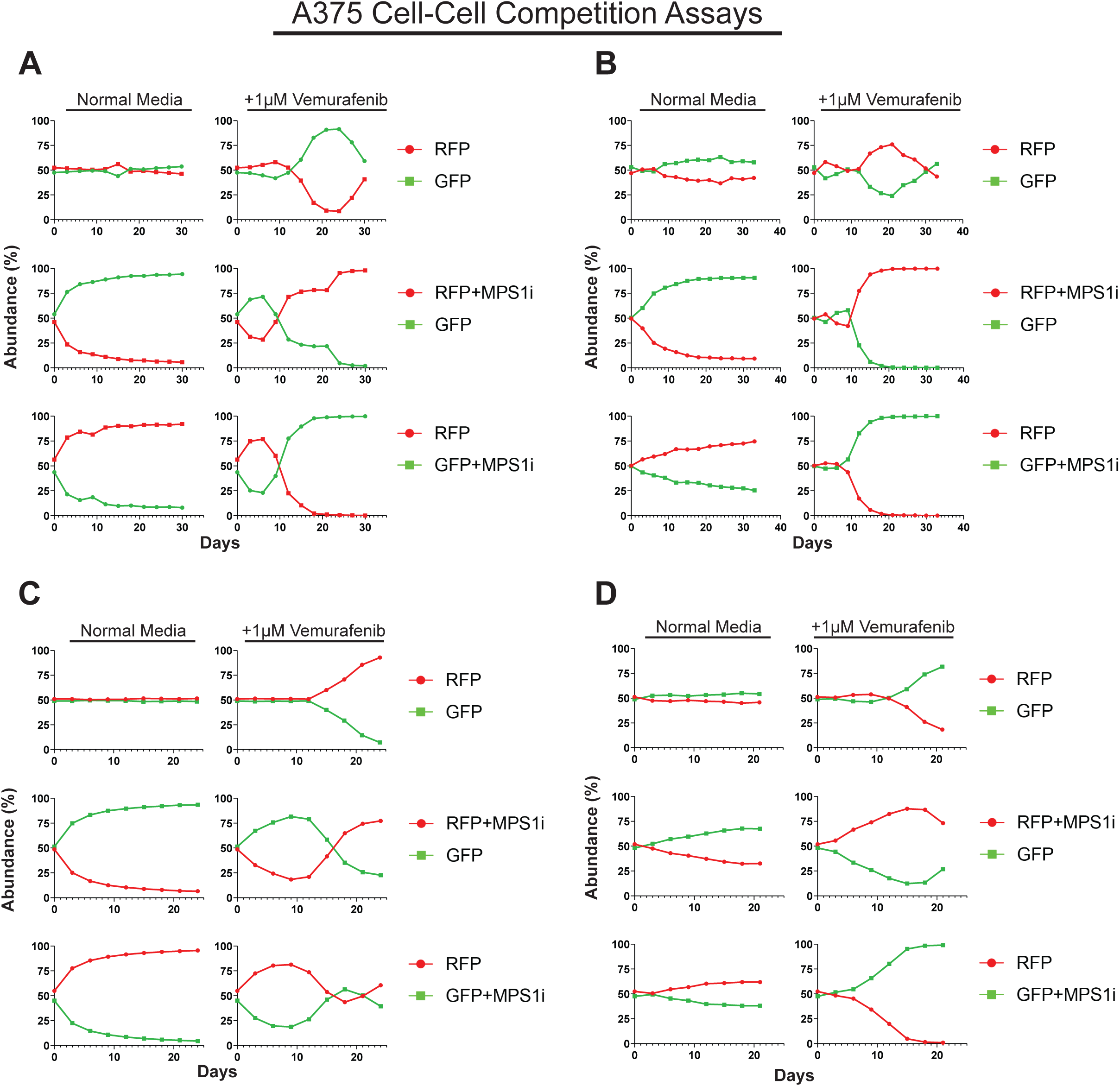
Transient exposure to an Mps1 inhibitor accelerates the acquisition of resistance to BRAF-inhibitors in multiple independent competitions. Related to Figure 1. (A-D) Relative abundance of competing A375 cell populations under normal growth conditions or vemurafenib treatment.

**Figure S3.**
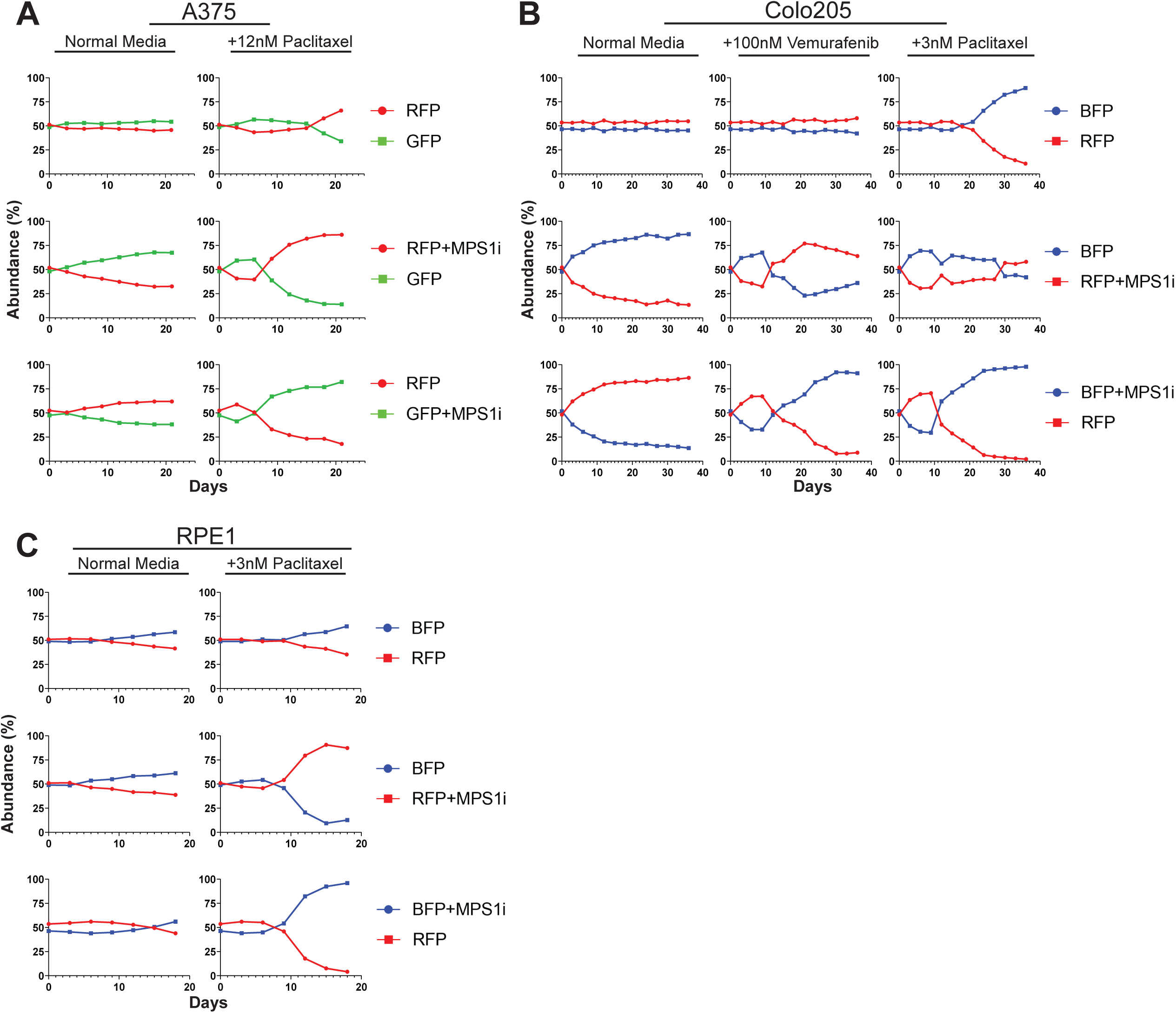
Transient exposure to an Mps1 inhibitor accelerates the acquisition of drug resistance in different genetic backgrounds in multiple independent competitions. Related to Figure 2. (A) Relative abundance of competing A375 cell populations under normal growth conditions or paclitaxel treatment. (B) Relative abundance of competing Colo205 cell populations under normal growth conditions, vemurafenib treatment, or paclitaxel treatment. (C) Relative abundance of competing RPE1 cell populations under normal growth conditions or varying levels of paclitaxel treatment.

**Figure S4.**
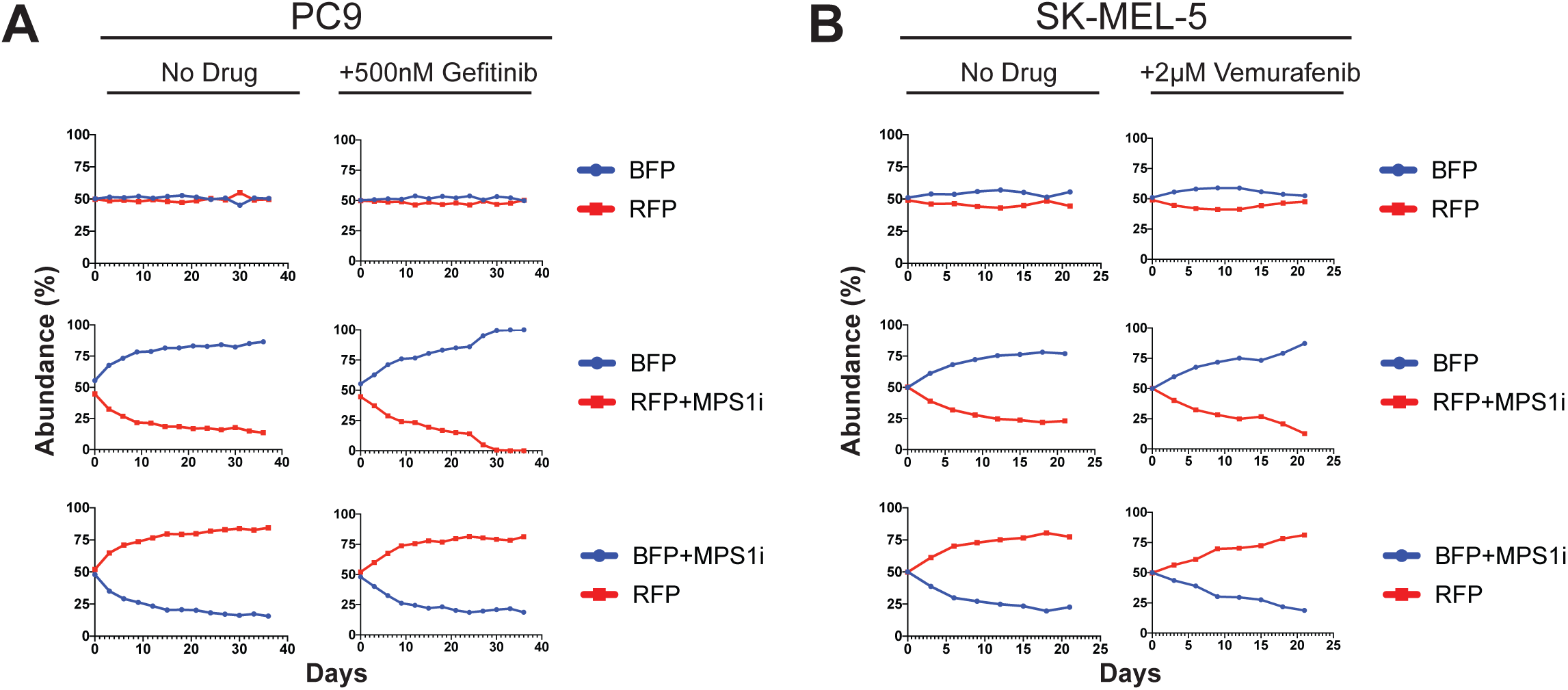
Transient Mps1 inhibition does not drive therapeutic resistance in all contexts. Related to Figure 2. (A) Relative abundance of competing PC-9 cell populations under normal growth conditions or gefitinib treatment. (B) Relative abundance of competing SKMEL-5 cell populations under normal growth conditions or vemurafenib treatment.

**Figure S5.**
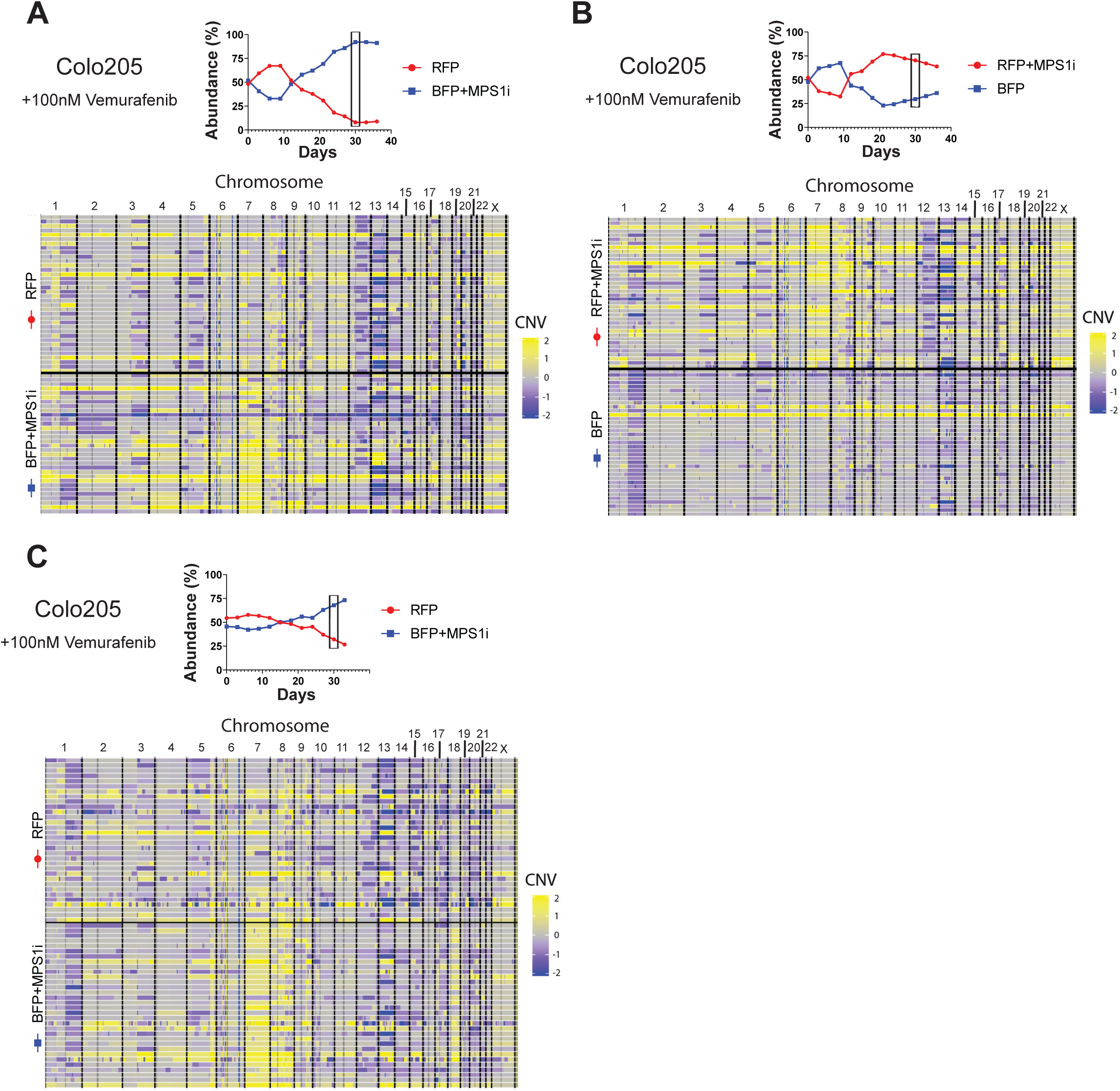
Single-cell sequencing identifies recurrent chromosomal changes in BRAFi-resistant populations of Colo205 cells. Related to Figure 3. (A) (Top) Plot displaying the competition of Colo205 cells in vemurafenib from which cells were isolated for single cell sequencing. Box indicates time point when cells were isolated. (Bottom) Heatmap displaying the karyotypic alterations of single cells isolated from the above competition. (B) (Top) Plot displaying the competition of Colo205 cells in vemurafenib from which cells were isolated for single cell sequencing. Box indicates time point when cells were isolated. (Bottom) Heatmap displaying the karyotypic alterations of single cells isolated from the above competition. (C) (Top) Plot displaying the competition of Colo205 cells in vemurafenib from which cells were isolated for single cell sequencing. Box indicates time point when cells were isolated. (Bottom) Heatmap displaying the karyotypic alterations of single cells isolated from the above competition. N.B. – Competition plots shown in (A), (B), and (C) were previously shown in Figures S3B, S3B, and 2B, respectively.

**Figure S6.**
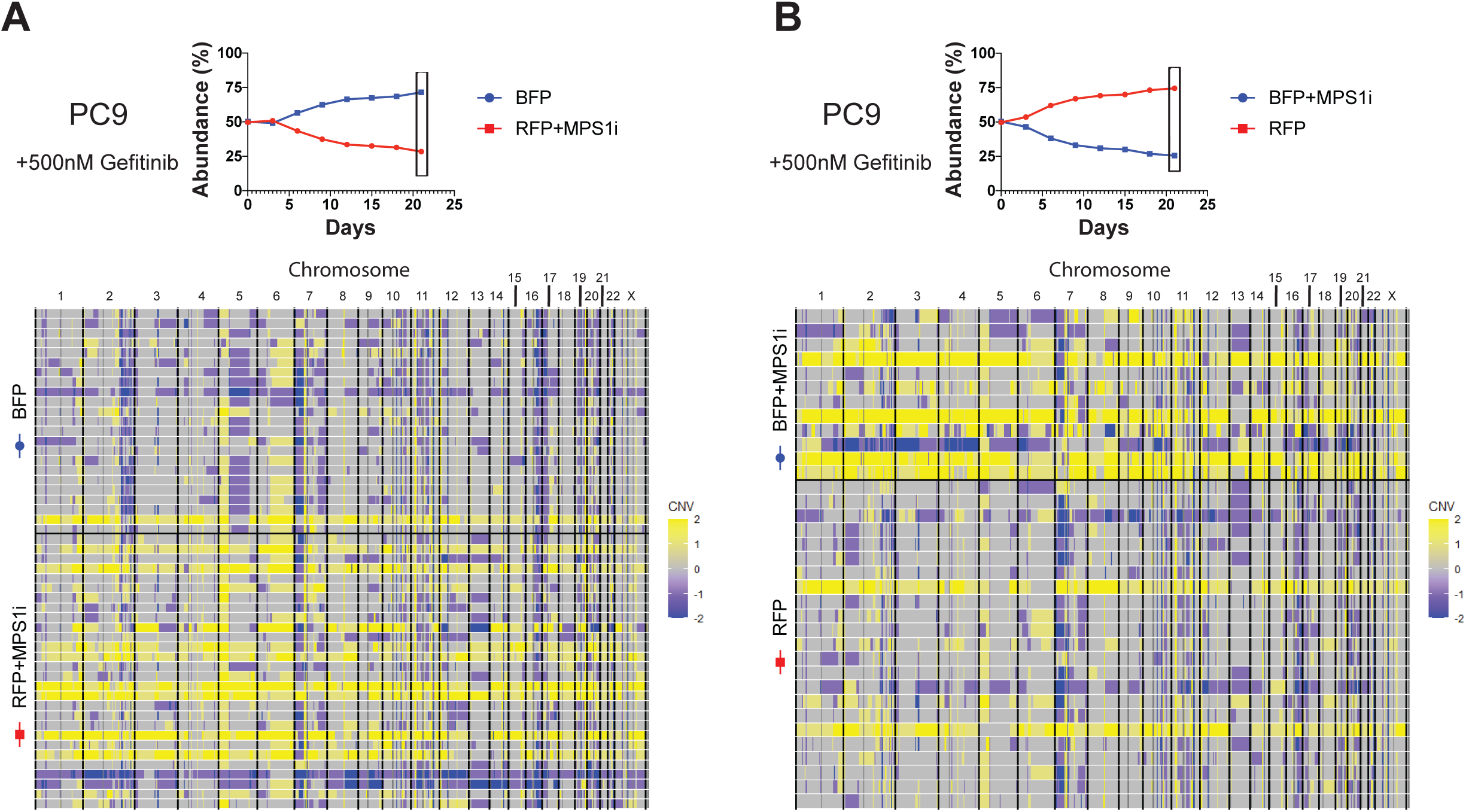
Single-cell sequencing identifies recurrent whole-genome duplications in Mps1i-treated PC9 cells. Related to Figure 3. (A) (Top) Plot displaying the competition of PC9 cells in gefitinib from which cells were isolated for single cell sequencing. Box indicates time point when cells were isolated. (Bottom) Heatmap displaying the karyotypic alterations of single cells isolated from the above competition. (B) (Top) Plot displaying the competition of PC9 cells in gefitinib from which cells were isolated for single cell sequencing. Box indicates time point when cells were isolated. (Bottom) Heatmap displaying the karyotypic alterations of single cells isolated from the above competition.

